# Heterologous caffeic acid biosynthesis in *Escherichia coli* is affected by choice of tyrosine ammonia lyase and redox partners for bacterial Cytochrome P450

**DOI:** 10.1101/707828

**Authors:** Kristina Haslinger, Kristala L.J. Prather

## Abstract

**Background:** Caffeic acid is industrially recognized for its antioxidant activity and therefore its potential to be used as an anti-inflammatory, anticancer, antiviral, antidiabetic and antidepressive agent. It is traditionally isolated from lignified plant material under energy-intensive and harsh chemical extraction conditions. However, over the last decade bottom-up biosynthesis approaches in microbial cell factories have been established, that have the potential to allow for a more tailored and sustainable production. One of these approaches has been implemented in *Escherichia coli* and only requires a two-step conversion of supplemented L-tyrosine by the actions of a tyrosine ammonia lyase and a bacterial Cytochrome P450 monooxygenase. Although the feeding of intermediates demonstrated the great potential of this combination of heterologous enzymes compared to others, no *de novo* synthesis of caffeic acid from glucose has been achieved utilizing the bacterial Cytochrome P450 thus far.

**Results:** The herein described work aimed at improving the efficiency of this two-step conversion in order to establish de novo caffeic acid formation from glucose. We implemented alternative tyrosine ammonia lyases that were reported to display superior substrate binding affinity and selectivity, and increased the efficiency of the Cytochrome P450 by altering the electron-donating redox system. With this strategy we were able to achieve final titers of more than 300 μM or 47 mg/L caffeic acid over 96 h in an otherwise wild type *E. coli* MG1655(DE3) strain with glucose as the only carbon source. We observed that the choice and gene dose of the redox system strongly influenced the Cytochrome P450 catalysis. In addition, we were successful in applying a tethering strategy that rendered even an initially unproductive Cytochrome P450/ redox system combination productive.

**Conclusions:** The caffeic acid titer achieved in this study is about 25% higher than titers reported for other heterologous caffeic acid pathways in wildtype *E. coli* without L-tyrosine supplementation. The tethering strategy applied to the Cytochrome P450 appears to be particularly useful for non-natural Cytochrome P450/redox partner combinations and could be useful for other recombinant pathways utilizing bacterial Cytochromes P450.

## Background

Caffeic acid is widely recognized for its medicinal potential due to its antidepressive [1], antihyperglycemic [2], anti-inflammatory [3], antioxidant [2, 4], anti-coagulatory [3], anticancer [5] and antiviral [6] properties. It is readily produced in plants as a key intermediate in phenylpropanoid biosynthesis. In this pathway, phenylalanine is diverted from primary metabolism by a phenylalanine ammonia lyase associated with the endoplasmatic reticulum and transformed into *trans*-cinnamic acid. Cinnamic acid is then hydroxylated by the membrane-anchored Cytochrome P450 enzymes cinnamate 4-hydroxylase (C4H) and *p*-coumarate 3-hydroxylase to *p*-coumarate and caffeic acid, respectively [7, 8]. From there a range of molecules can be produced that serve as lignin building blocks or precursors for secondary metabolites such as tannins, (iso)flavonoids, anthocyanins, stilbenes and coumarins [9]. All of these compounds have high market value but are difficult to isolate because they are of low natural abundance (e.g. stilbenes and coumarins), or challenging to extract (e.g. lignin-derived aromatics) [10]. Therefore, over the last decade various strategies have been developed to implement biosynthetic pathways in microbial cell factories that promise their tailored biosynthesis in a sustainable manner. Recent examples are the production of stilbenoids and flavonoids in *Corynebacterium glutamicum* [11, 12], and curcumin [13, 14] and caffeic acid [14–24] in *Escherichia coli*. For the biosynthesis of *p*-coumaric acid in *E. coli*, it was found that using L-tyrosine as a pathway precursor was superior over phenylalanine [25], since activity of the plant Cytochrome P450 enzyme C4H could not be reconstituted as of recently [26]. Based on this finding, two major strategies have been devised to produce caffeic acid that employ microbial tyrosine ammonia lyases (TAL) to generate *p*-coumaric acid followed by either (1) a flavin-dependent HpaBC-type oxidoreductase complex (4-hydroxyphenylacetate 3-hydroxylase, PFAM PF03241) from *Saccharothrix espanaensis* [14–18], *E. coli* [19–21], *Thermus thermophilus* HB8 [20] or *Pseudomonas aeruginosa* [22, 23], or (2) a bacterial cytochrome P450 enzyme CYP199A2 F185L from *Rhodopseudomonas palustris* [14, 18, 24]. In all of these studies it became evident that the caffeic titers are rather low unless l-tyrosine or *p*-coumaric acid are added to the growth media, or the aromatic amino acid pathway is engineered to increase intracellular L-tyrosine levels. For the pathways utilizing HpaBC-type oxidoreductases, the highest titer reported for *de novo* synthesis in wild-type *E. coli* to date is 42 mg/L (*S. espanaensis* TAL and HpaBC) [17]. However, to our knowledge no *de novo* synthesis has been reported for pathways utilizing CYP199A2 F185L.

In this study, we established *de novo* biosynthesis of caffeic acid from glucose through the actions of TAL and CYP199A2 F185L NΔ7. In order to achieve this goal, we tested TALs from three different organisms and explored strategies to enhance the activity of CYP199A2 F185L NΔ7. We found that driving the binding equilibrium of the electron-donating redox partners to CYP199A2 F185L NΔ7 towards the bound state improves pathway titers and enabled us to produce ~47 mg/L caffeic acid from glucose in wildtype *E. coli* MG1655(DE3). This titer is slightly higher than the titers reported for the HpaBC-based pathways in wildtype *E. coli* with glucose as the only carbon source [17, 19].

## Results

In an earlier study Rodrigues et al. demonstrated the two-step conversion of 3 mM l-tyrosine to caffeic acid in *E. coli* MG1655(DE3) expressing the enzymes RgTAL and CYP199A2 F185L NΔ7 with redox partners, without reporting *de novo* production of caffeic acid from glucose (Figure 1) [18]. In this study we set out to improve these enzymatic steps to establish caffeic acid production from glucose without supplementing L-tyrosine. When examining the two-step conversion more closely, we determined that both pathway steps needed improvement. First, the efficiency of the committed step, the conversion of L-tyrosine to *p*-coumaric acid, determines how much L-tyrosine is withdrawn from primary metabolism and fed into the pathway. Therefore, we hypothesized that TAL variants with higher selectivity and affinity for L-tyrosine would improve pathway flux. Second, the hydroxylation of *p*-coumaric acid to caffeic acid catalyzed by CYP199A2 F185L NΔ7 appears to be a bottleneck in the pathway, since *p*-coumaric acid accumulates in the fermentation [18]. This accumulation is thought to be detrimental because *p*-coumaric acid has been shown to inhibit TAL activity and to be cytotoxic [27, 28]. Since a common problem with Cytochrome P450 catalyzed reactions is the protein-protein interaction with redox partners, which is strictly required for electron transfer and substrate turnover [29], we hypothesized that driving the assembly of the redox complex would lead to higher product titers.

**Figure 1:**
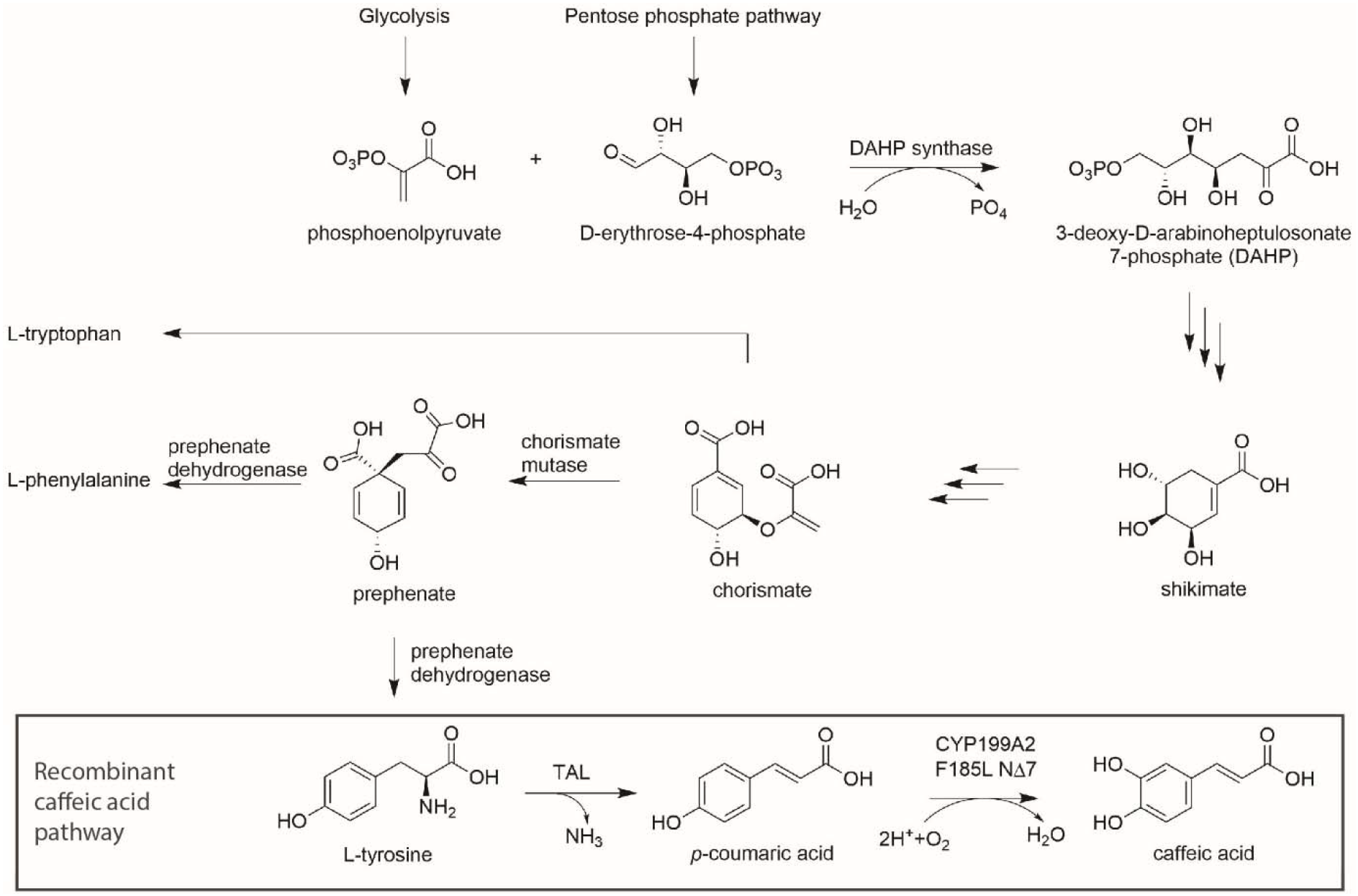
Aromatic amino acid anabolism and recombinant caffeic acid pathway with l-tyrosine as a branchpoint, and TAL and CYP199A2 F185L N□7 catalyzing the two pathway steps.

To improve the first pathway step, we selected two homologous tyrosine ammonia lyases with supposedly superior characteristics compared to RgTAL, namely a stronger selectivity for L-tyrosine over L-phenylalanine, higher substrate affinity (Km) and superior catalytic efficiency (k_cat_/K_m_) (Supporting Information Table S1) [30]. We chose FjTAL from *Flavobacterium johnsoniae* and SeSam8 from *Saccharothrix espanaensis* and obtained the synthetic genes codon-optimized for expression in *E. coli*. In a first pass, utilizing these two TALs in the same three plasmid expression system as used by Rodrigues et al., and providing glucose as the only carbon source, we observed accumulation of caffeic acid 72 h post induction (p. i.). The highest titers of caffeic acid and *p*-coumaric acid are seen with the FjTAL enzyme (Figure 2A, strain s02). In a parallel experiment, where 3 mM L-tyrosine was fed in addition to glucose, the final caffeic acid titers were comparable among the three strains (Figure 2B). This indicates that all enzymes are able to efficiently route L-tyrosine into the caffeic acid pathway at high L-tyrosine concentrations, whereas FjTAL outperforms the other enzymes under low L-tyrosine conditions and is therefore a strong candidate for this pathway.

**Figure 2:**
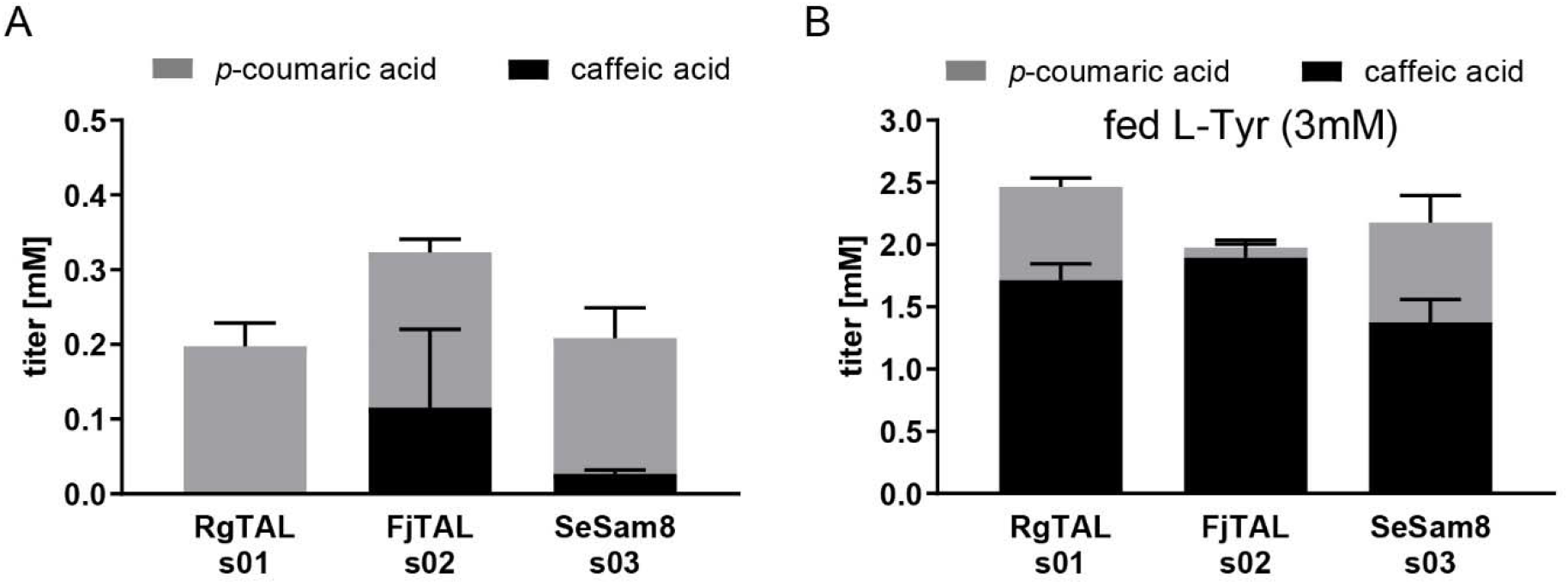
Titers of *p*-coumaric acid and caffeic acid from glucose without (A) and with (B) L-Tyr supplementation (stacked histograms, error bars= standard deviation of biological replicates, n≥3).

Next, we sought to improve the efficiency of the second pathway step, the hydroxylation of *p*-coumaric acid to caffeic acid catalyzed by CYP199A2 F185L NΔ7, by enhancing the efficiency of the electron transfer step from the two redox partner proteins to CYP199A2 F185L NΔ7. To achieve this goal, we tested three strategies: 1. the use of alternative redox partners, 2. the tethering of the redox complex by creating genetic fusions with high-affinity tethering domains, and 3. the supply of extra gene copies coding for one of the redox partners. To facilitate cloning from here on in the study, we use both multiple cloning sites of the pETDuet vector for the genes encoding redox enzymes rather than the bicistronic pKVS45 vector (see Tables 1 and 2).

**Table 1:**
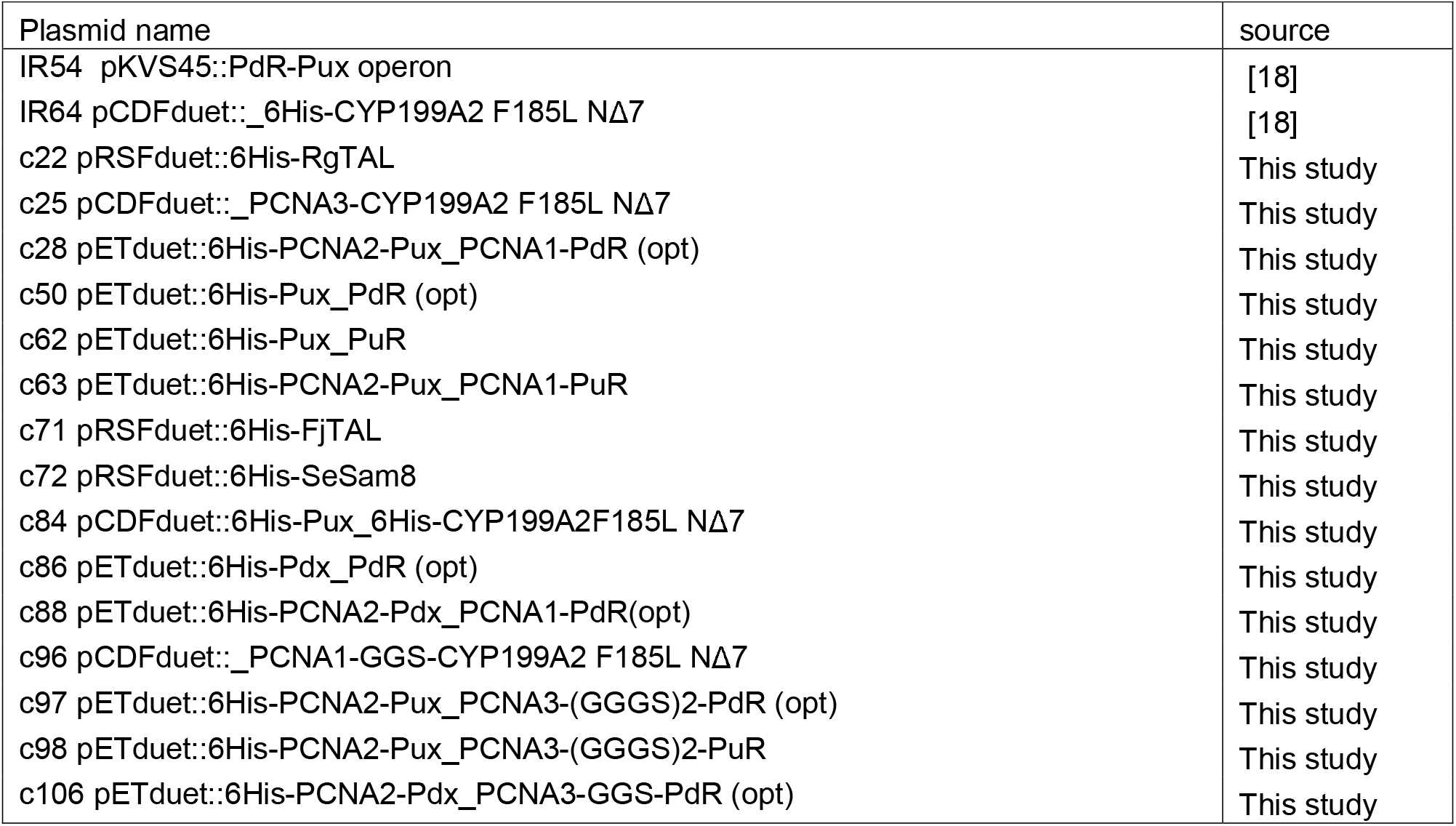
List of plasmids used in caffeic acid production strains.

**Table 2:**
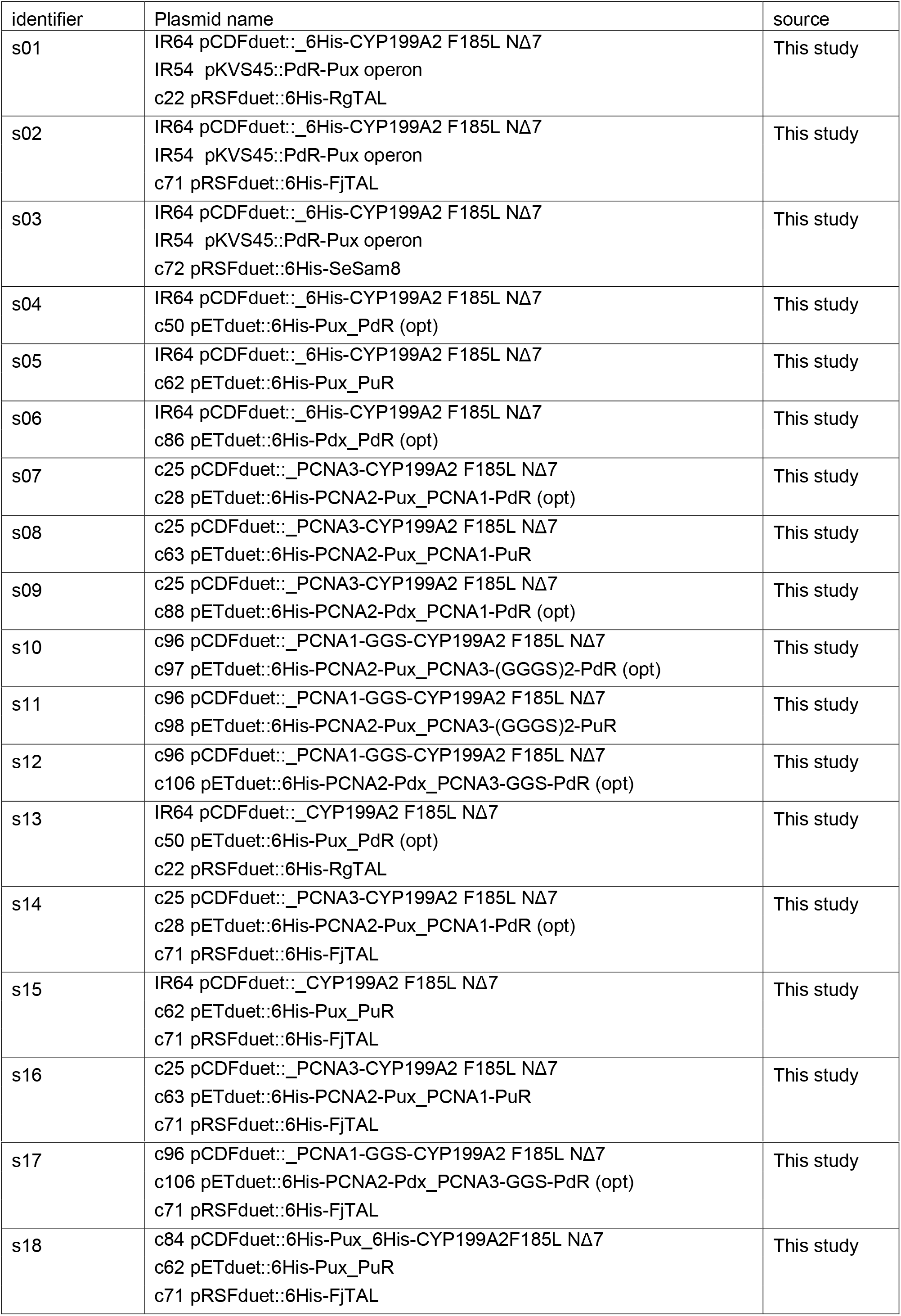
List of *E. coli* MG1655(DE3) strains used in fermentation experiments.

For class I Cytochromes P450, two redox partners are required to provide two electrons from NAD(P)H: an iron-sulfur cluster containing ferredoxin (Fdx) and a flavin-dependent ferredoxin reductase (FdR) [31]. Rodrigues et al. utilized a redox system composed of palustrisredoxin (Pux) and putidaredoxin reductase (PdR), which had been used in the original characterization of CYP199A2 [32]. This is, however, not the natural redox system for CYP199A2, since the palustrisredoxin reductase PuR was only identified and characterized a few years later [33]. Although the Pux/PdR redox system has been proven to support substrate turnover, it remained unclear whether the assembly of the trimeric complex and the respective redox potentials of the proteins supported optimal electron transfer. Therefore, we decided to test the natural redox system (Pux/PuR) alongside a well-characterized surrogate redox system (Pdx/PdR). We determined the caffeic acid titers 72 h p. i. with supplementation of *p*-coumaric acid for three strains expressing CYP199A2 F185L NΔ7 and one of the three respective redox systems Pux/PdR (hybrid, s04), Pux/PuR (natural, s05), Pdx/PdR (surrogate, s06). We observed the highest titers for the natural redox system (s05) and no turnover with the full surrogate system composed of Pdx/PdR (Figure 3A). This suggests that the electron transfer from ferredoxin to CYP199A2 F185L NΔ7 is severely impaired with the surrogate ferredoxin Pdx, whereas the electron transfer from PdR to Pux in the hybrid system appears to sufficiently support substrate turnover. The native redox complex Pux/PuR, however, displays the highest catalytic power and a titer of 1.6 +/- 0.32 mM caffeic acid was observed which corresponds to 53% conversion of the fed *p*-coumaric acid. These results indicate that the careful choice of redox system is crucial for this pathway step.

**Figure 3:**
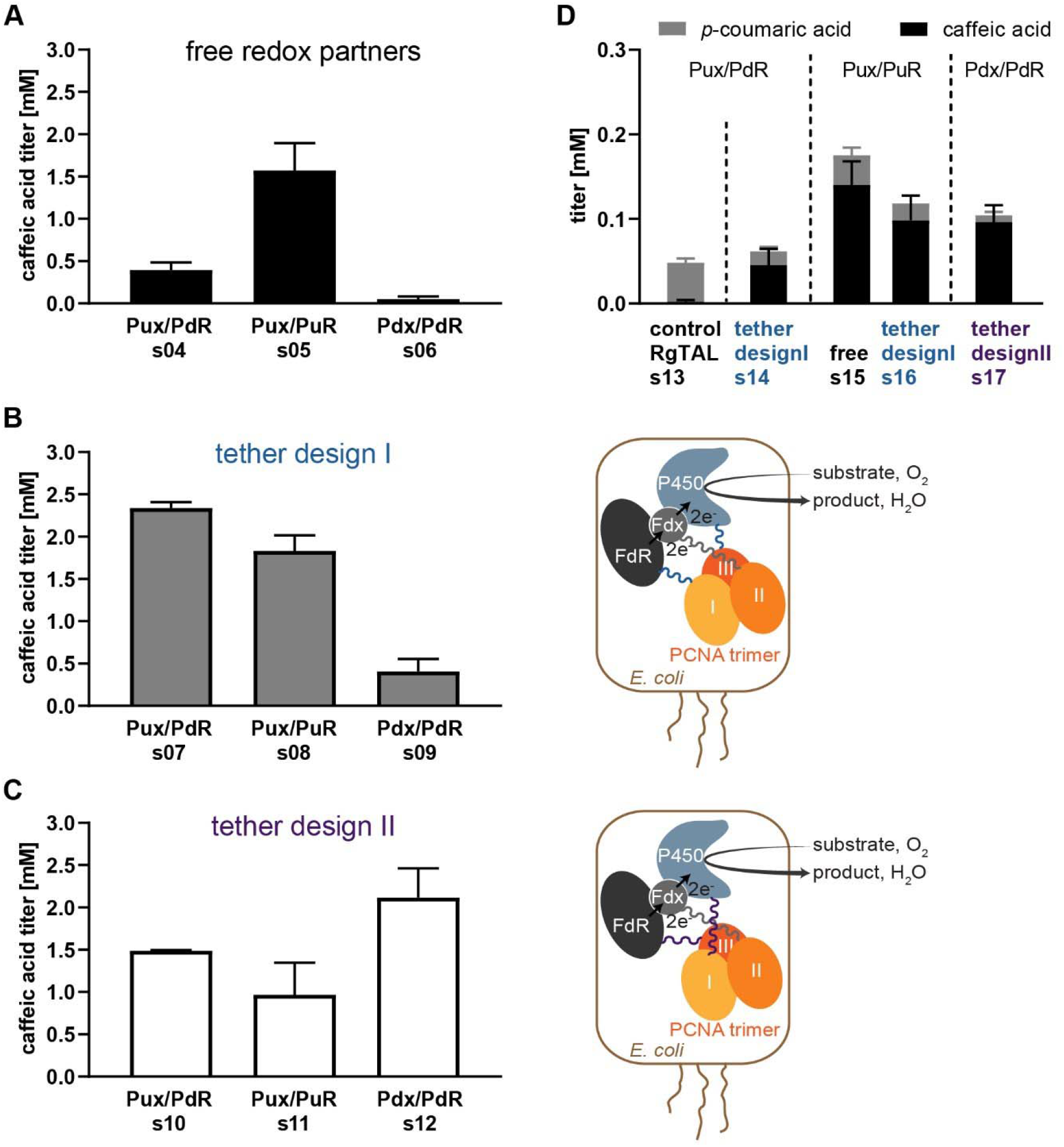
The choice of redox partners and tethering strategies for redox partners leads to higher caffeic acid titers from *p*-coumaric acid (panels A-C) and from glucose (panel D). A-C, caffeic acid titers from 3 mM *p*-coumaric acid 72 h p. i.: untethered/free redox partners (A), tether design I analogous to PUPPET [35] (B), tether design II (C). Stacked histograms of *p*-coumaric and caffeic acid titers after 72 h of fermentation for select strains expressing the two-step pathway (D). (Error bars= standard deviation of biological replicates, n?3; Pictograms of tether designs: P450=Cytochrome P450 enzyme, Fdx=ferredoxin (Pux or Pdx), FdR=ferredoxin reductase (PuR or PdR)).

With our second strategy, we sought to further improve these redox systems by generating genetic fusions of the enzymes with the subunits of the heterotrimeric DNA sliding clamp PCNA (Proliferating Cell Nuclear Antigen) of *Sulfolobus solfataricus* P2 [34]. This PCNA complex has been shown to tolerate the fusion of other genes to the ‘3 ends (C-termini) [35] of its three subunits, while maintaining their high binding affinity towards each other: the PCNA1/PCNA2 dimer has a dissociation constant in the low picomolar range and the PCNA1/PCNA2/PCNA3 trimer in the high nanomolar range [34]. This fusion strategy has been shown to be highly efficient for the *in vitro* reconstitution of Cytochrome P450 activity and was termed PUPPET by the inventors (<u>P</u>CNA-<u>u</u>tilized <u>p</u>rotein <u>c</u>omplex of <u>P</u>450 and its two <u>e</u>lectron <u>t</u>ransfer-related proteins) [35–40]. To our knowledge, this strategy hasn’t been used in whole-cell catalysis to date. Initially, we tested fusion proteins analogous to the previously described PUPPET fusions with FdR fused to the C-terminus of PCNA domain 1, Fdx to PCNA2 and the Cytochrome P450 to PCNA3 (tether design I, strains s07-s09; Figure 3B). When feeding 3 mM *p*-coumaric acid, we observed higher titers of caffeic acid for all tethered redox systems than compared to the respective free enzymes. The effect was more pronounced with the hybrid and surrogate systems, where a 6-fold increase in titer was observed for Pux/PdR (s07) and an 8-fold increase for Pdx/PdR (s09). Overall, the highest titer was observed with the tethered version of Pux/PdR (s07, titer: 2.3 +/- 0.07 mM). Next, we investigated whether these titers could be further improved by generating a new arrangement of the fusion partners. Based on the published dissociation constants for the well-studied Cytochrome P450 CYP101A1 and its redox partners [41, 42], we assumed that the affinity of Fdx to FdR is about 100-fold higher than the affinity of Fdx to the Cytochrome P450. We hypothesized that the high affinity interaction between PCNA1 and PCNA2 might be even more beneficial to the low affinity interaction between the Cytochrome P450 and Fdx than between Fdx and FdR. Therefore, we generated a second set of fusion genes (tether design II), where CYP199A2 F185L NΔ7 is fused to PCNA1, Fdx to PCNA2 and FdR to PCNA3, while maintaining the linker arrangements that had previously been optimized for the respective elements of the redox complex [39] (s 10-s12, Fig 3C). With these alternative tethering constructs, the highest final caffeic acid titers were obtained with the surrogate Pdx/PdR redox system (s10, titer: 2.1 +/- 0.35 mM), while the titers obtained with the other redox systems were lower than in the previous experiments. This indicates that the domain arrangements in the second tether design supports the weaker protein-protein interactions in the surrogate redox complex better than the other tether design, whereas it disturbs catalysis with the two redox systems that already led to high titers with free redox partners and tether design I.

Next, we tested the best redox partner constructs in the context of the full pathway with FjTAL as the first pathway enzyme (Figure 3D). We observed the highest caffeic acid titers with the untethered, natural redox partners (Pux/PuR, s15, titer: 0.14 +/0.028 mM). Although strains s07, s08 and s12 had slightly outperformed s05 in the one-step conversion, the corresponding strains expressing FjTAL (s14, s16, s17, respectively) yielded lower caffeic acid titers in the two-step recombinant pathway. The cost for expressing the additional tethering domains may offset the positive effects of the enhanced enzymatic activity. In all of the fermentations, lower final titers of *p*-coumaric acid are measured than in the initial test of FjTAL (s02), which indicates that the changes made to the second pathway step allow for an almost complete conversion to the final product.

Lastly, we tested whether additional copies of the palustrisredoxin encoding gene, *pux*, would further improve the performance of the so far best pathway configuration with FjTAL and the natural redox partners of CYP199A2 F185L NΔ7 (Pux/PuR redox system). Therefore, we inserted *pux* into MCS1 of plasmid IR64 pCDFDuet::_6His-CYP199A2 F185L NΔ7, yielding plasmid c84 pCDFDuet::6His-Pux_6His-CYP199A2F185L NΔ7. Based on the supplier’s reports (Novagen), the copy numbers of pETDuet and pCDFDuet are in a similar range so that the incorporation of an additional gene copy into pCDFDuet should lead to an estimated doubling of the gene dose and potentially the level of protein expressed. When comparing the strain harboring this set of plasmids (s18) to the RgTAL control strain (s13) and the strain expressing FjTAL and Pux/PuR (s15), we observed an increase in caffeic acid titer with full consumption of the intermediate *p*-coumaric acid (Figure 4A). This indicates that the availability of Pux was previously insufficient and that a higher expression level of this protein supports better Cytochrome P450 performance. Despite the improvements in final caffeic acid titer, we observed an accumulation of *p*-coumaric acid in early fermentation until 48 h p. i. and then a sharp drop in titer until it is fully converted to caffeic acid at 96 h p. i. (Figure 4B). This indicates that in early fermentation the first pathway step is still faster than the second pathway step. In late fermentation, the conversion of *p*-coumaric acid to caffeic acid is faster than the formation of the intermediate, or no additional *p*-coumaric acid is formed. This could be caused by the lack of available L-tyrosine once the cultures reach stationary phase, although we did not observe increased titers when spiking the cultures with 3 mM l-tyrosine at 48 h p. i. (I Figure S1). Therefore, we are inclined to suggest that the tyrosine ammonia lyase has lost its activity by that time. Potential causes could be structural instability of the TAL enzyme or its inhibition by the intermediate as described previously [27]. Overall, with the exchange of RgTAL for FjTAL and the change of the redox system from Pux/PdR to Pux/PuR with an additional gene copy of *pux*, we improved this recombinant pathway and were able to produce caffeic acid from glucose without feeding L-tyrosine. The highest final titer after 96 h of fermentation was 47 mg/L, which is slightly higher than caffeic acid titers achieved with other recombinant pathways without L-tyrosine supplementation [17, 19]. Furthermore, the improved pathway is able to convert >50% of fed L-tyrosine to caffeic acid (SI Figure S1), which indicates that it should be able to produce high amounts of caffeic acid in a tyrosine-producer strain.

**Figure 4:**
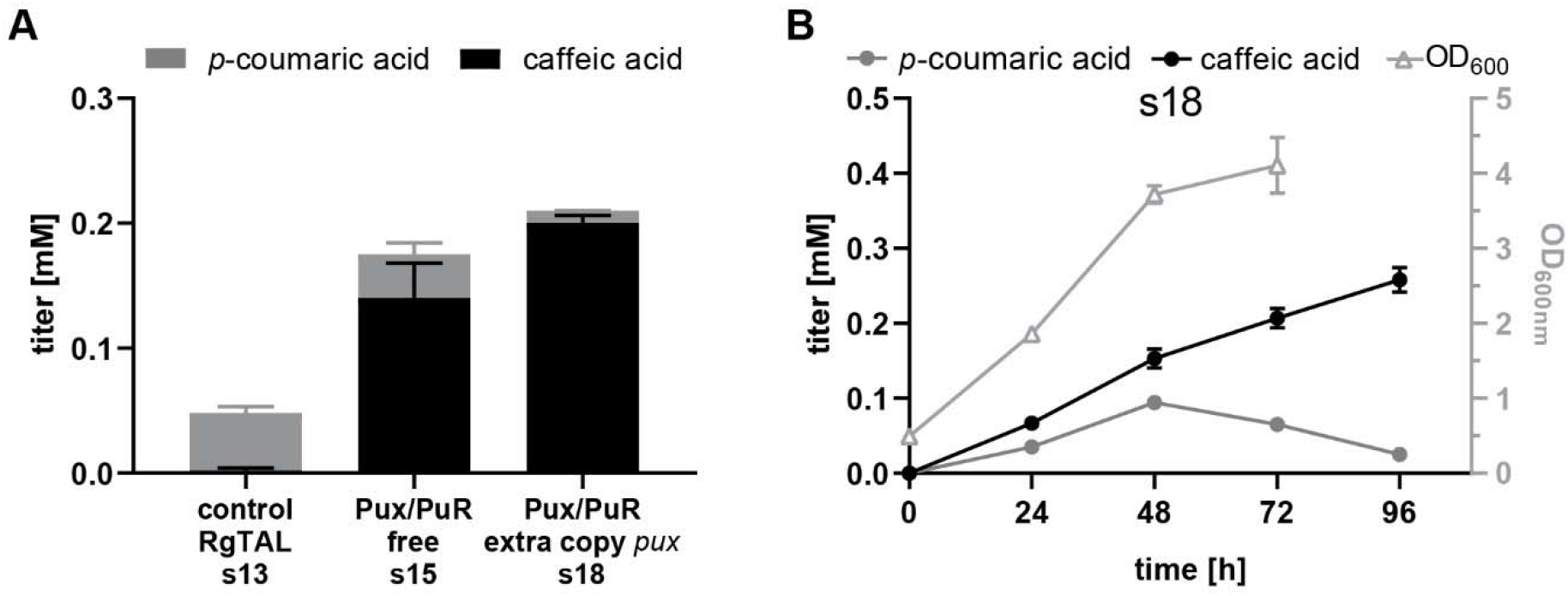
Duplication of the *pux* gene copy number further increases caffeic acid titers. Stacked histograms of *p*-coumaric and caffeic acid titers after 72 h of fermentation with glucose as the only carbon source for select strains expressing the two-step pathway (A). Titers plotted over time of a 96 h fermentation of s18 (B). (Error bars= standard deviation of biological replicates, n≥3.)

## Discussion

Building microbial cell factories for the production of plant polyphenols has been a major goal for metabolic engineers over the last decade [43, 44]. The low abundance of these compounds and their occurrence in complex mixtures of variable composition in plants, makes recombinant microbial cell factories an attractive source for industrial applications. However, the strict regulation of the aromatic amino acid metabolism, which provides precursors to most recombinant polyphenol-producing pathways, limits the overall pathway efficiency. For recombinant polyphenol-producing pathways in *E.coli*, it has been observed that overcoming the precursor bottleneck by metabolic engineering of the aromatic amino acid pathway, often reveals bottlenecks further down the recombinant pathway [45–47]. Therefore, it is crucial to optimize the recombinant pathway itself before moving into a microbial chassis with deregulated aromatic amino acid production. In this study, we optimized the two-step conversion of L-tyrosine to caffeic acid. Here it is important to ensure high efficiency of the second pathway step to avoid accumulation of *p*-coumaric acid, which has been shown to severely inhibit the activity of the first pathway enzyme, TAL [27]. The three strategies we tested focused on the electron-donating redox partners rather than the Cytochrome P450 enzyme itself. Previous *in vitro* studies of this particular Cytochrome P450 and others have shown that the right choice of redox system, in particular the ferredoxin, is crucial for efficient electron transfer and enzyme catalysis [29, 33]. As expected, we observed the highest caffeic acid titers with the natural redox system composed of Pux and PuR in the one-step conversion with untethered redox partners. However, when we applied tethering strategies to increase the affinity of the Cytochrome P450 and the redox partners towards each other, we observed higher titers with the non-natural redox partners. Tethering strategies have previously been applied to several Cytochrome P450 enzymes, both *in vitro* [35, 42, 48–51] and *in vivo* [42, 48]. The *in vitro* studies showed that tethered redox complexes are able to overcome the need to use an excess of redox partners over the Cytochrome P450 enzyme, to compensate for low protein-protein affinities (typically a five-to twenty-fold molar excess of ferredoxin is used *in vitro*). Furthermore, kinetic studies showed that at low enzyme concentrations, the tethered complexes outperform the 1:1:1 mixtures of free enzymes. These reports and our findings for our versions of the PUPPET tether indicate that tethering strategies in whole-cell catalysis may be particularly useful in two scenarios: (A) if the expression levels of the Cytochrome P450 and redox partners are low (poor protein expression, expression from genomic gene copies or as part of a multi-enzyme recombinant pathway), or (B) if the natural redox partners are unknown and surrogate systems are used to reconstitute the Cytochrome P450 activity.

To our knowledge, this study is the first one to use the PUPPET tether in whole-cell catalysis and also the first one to use any of the known Cytochrome P450 tethers in the context of a recombinant pathway. In the caffeic acid pathway, the tethered Cytochrome P450 complexes were slightly outperformed by the free, natural redox complex, in particular in the presence of extra copies of the *pux* gene (s18). This may indicate that the metabolic burden of expressing the PCNA subunits in addition to the pathway enzymes and the three resistance genes required for plasmid maintenance represents a disadvantage of the strains expressing the tethered Cytochrome P450 complexes compared to the ones expressing the free, natural redox complex (s15 and s18). The fact that s18 outperforms s15 indicates that the availability of Pux is limiting in s15, and is in good agreement with observations made in other whole-cell conversions [52, 53]. Since our strategy only doubled the gene dose of *pux*, it is very well possible that rearranging the genes in the vector system to achieve higher protein levels of Pux relative to the other enzymes could lead to even better results than described in this study. Our optimization efforts of the second pathway step in combination with the use of FjTAL for the first pathway step, enabled us to demonstrate the *de novo* production of caffeic acid in an otherwise wild type *E. coli* background. FjTAL had previously been seen to be beneficial for the production of *p*-coumaric acid and its derivatives in other microbes [11, 54, 55], however, to our knowledge it has not been used in *E. coli*. It appears that this enzyme allows for a more efficient routing of L-tyrosine into the caffeic acid pathway than RgTAL at low L-tyrosine concentrations. Under high L-tyrosine conditions, at levels that we would expect in tyrosine producer strains [56], our fermentation strains expressing FjTAL achieve slightly higher caffeic acid titers than the strains expressing RgTAL and lower titers of *p*-coumaric acid. This indicates that the optimized pathway is more balanced so that less *p*-coumaric acid accumulates but overall less L-tyrosine is converted into *p*-coumaric acid. To further improve these results, it is necessary to investigate the stability and activity of the FjTAL enzyme over time, since it appears to be inactive after 48h of fermentation.

## Conclusions

In this study we established *de novo* synthesis of caffeic acid by expressing tyrosine ammonia lyase from *Flavobacterium johnsoniae* and CYP199A2 F185L N 7 from *Rhodopseudomonas palustris* with its redox partners palustrisredoxin and palustrisredoxin reductase. We found that compared to earlier versions of this pathway, changes made to the redox partners, namely the use of palustrisredoxin reductase instead of putidaredoxin reductase and the duplication of the palustrisredoxin gene dose, as well as the use of FjTAL instead of RgTAL, enhanced the pathway performance under low L-tyrosine conditions as encountered in otherwise wild type *E. coli*. Furthermore, we observed that applying a tethering strategy to the Cytochrome P450-catalyzed pathway step based on the PUPPET system [35] increases caffeic acid titers in strains expressing non-natural redox systems. This indicates that this strategy can be useful for pathways containing orphan bacterial Cytochromes P450. The optimized caffeic acid pathway could now be transferred into a tyrosine-producer *E. coli* strain for more in-depth characterization or process engineering.

## Materials and Methods

### Bacterial strains and plasmids

All molecular cloning and plasmid propagation steps were performed in chemically competent *Escherichia coli E. cloni*^®^ 10G (F-*mcrA* Δ(*mrr-hsd*RMS-*mcr*BC) *endA1 recA1* Φ80*dlac*ZΔM15 Δ*lac*X74 *ara*D139 Δ(*ara,leu*)7697*gal*U *gal*K *rps*L *nup*G λ-*ton*A) produced by Lucigen (Middleton, WI, USA). Gene expression under the control of T7 promoters was performed in *E. coli* K-12 MG1655(DE3) [57]. Plasmids were constructed with a range of strategies summarized in the Supplementary Information (SI) Table S2. All genes in the final constructs were fully sequenced (Eton Bioscience, Charlestown, MA). The FjTAL, SeSam8 and PCNA1-PdR genes were codon optimized for *E. coli* and synthesized as gblocks^®^ gene fragments by Integrated DNA Technologies (Coralville, IA, USA) (sequence provided in SI). Plasmids pHSG-PCNA2 and pHSG-PCNA3 were a gift from Teruyuki Nagamune obtained through Addgene (Cambridge, MA, USA) (Addgene plasmid # 66126; http://n2t.net/addgene:66126; RRID:Addgene_66126) and (Addgene plasmid # 66127; http://n2t.net/addgene:66127; RRID:Addgene_66127) [35]. Plasmid pACYCDuet-PuR/Pux was a gift from Dr. Stephen G. Bell (University of Adelaide, Australia). The construction of plasmids IR54 and IR64 is described in Rodrigues et al 2015 [18].

The peptide linkers connecting the PCNA subunits with the respective enzymes were designed based on the optimized linkers described in Haga et al. 2013 [39] (tether design I: PCNA1-(GGGS)_2_-FdR, PCNA2-GGGSP_20_G-Fdx, PCNA3-GGS-Cytochrome P450; tether design II: PCNA1-GGS-Cytochrome P450, PCNA2-GGGSP_20_G-Fdx, PCNA3-(GGGS)_2_-FdR).

### Fermentation

Plasmids and strains used in fermentations are described in Table 1 and Table 2, respectively. *E. coli* K-12 MG1655(DE3) made chemically competent according to the protocol by Inoue et al [58] was sequentially transformed with appropriate plasmids. The correct identity of strains was confirmed by colony PCR. Starter cultures were prepared from three individual colonies of the final strains in 5 mL Lysogeny broth (LB) supplemented with carbenicillin (100 μg/mL), spectinomycin (50 μg/mL) and kanamycin (50 μg/mL, only s01-s03 and s13-s18) in round-bottom polystyrene tubes, incubated over night at 37°C with agitation and used to inoculate the main cultures (7 mL LB with antibiotics; round-bottom polystyrene tubes). After 4 h of growth at 37°C, 250 rpm, OD_600_ was measured and the appropriate volume of each culture pelleted and resuspended in modified, selective M9 including substrates and 4% glucose to obtain 15 mL cultures at OD_600_ of 0.7 or 20 mL cultures at OD_600_ of 0.5 to 0.7 (time course experiment) in sterile glass tubes. These cultures were incubated at 26°C, 160 rpm for 72 h or 96 h (time course experiment). For the time course experiment samples of 1000 μL were taken every 24 h, for all other experiments samples of 2000 μL were taken after 72 h and either stored at -20°C until further processing or extracted with ethyl acetate immediately.

M9 medium composition (1x) prepared from sterile stocks: M9 salts (Millipore-Sigma, used as 5x stock), Trace Mineral Supplement (ATCC^®^ MD-TMS^™^, used as 200x stock), vitamin mix (from 100x stock; final: riboflavin 0.84 mg/L, folic acid 0.084 mg/L, nicotinic acid 12.2 mg/L, pyridoxine 2.8 mg/L, and pantothenic acid 10.8 mg/L), biotin (from 1000x stock; final: 0.24 mg/L), thiamine (from 1470x stock; final: 340 mg/L), δ-Aminolevulinic acid (from 1000x stock in MeOH, final: 7.5 μg/mL), IPTG (from 1000x stock, final: 1 mM), aTc (from 1000x stock, final: 100 ng/mL; only included in fermentations of s01-s03), carbenicillin (from 1000x stock, final: 100 μg/mL), spectinomycin (from 1000x stock, final: 50 μg/mL), kanamycin (from 1000x stock, final: 50 μg/mL, only strains s01-s03 and s13-s18), 4% (w/v) glucose (from 50% w/v stock). Optional: *p*-coumaric acid (from fresh 100x stock in MeOH, final 3 mM) or L-tyrosine (from fresh 100x stock in 1M HCl).

### Product extraction

The samples were acidified with 6N HCl (pH<3) and split into two tubes as technical duplicates. Samples were extracted twice with equal volumes of ethylacetate. The organic phases of both extraction steps were combined and evaporated under a stream of air or nitrogen. The dried material was resuspended in 100 μL Acetonitrile with 0.1% Trifluoracetic acid (10x concentrated compared to culture) or 80 μL Acetonitrile with 0.1% Trifluoracetic acid (5x concentrated compared to culture) for the time course experiment. Samples were transferred into HPLC vials with conical glass inserts and analyzed by HPLC.

### HLPC analysis

10 μL of the samples were analyzed by reversed-phase HPLC (instrument: Agilent 1100, column: Agilent Zorbax Eclipse XDB-C18 80Å, 4.6 x 150 mm, 5μm; detector: Agilent diode array detector G1315B, λ=310nm, gradient: 10% to 20% Acetonitrile with 0.1% Trifluoracetic acid over 17 min. The *p*-coumaric acid and caffeic acid peaks were identified by comparing the retention times to authentic standards and by mass spectrometry (Agilent G6120, quadrupole MS). The integrated peak areas were converted to concentrations in mM based on calibration curves generated with authentic standards.

## List of abbreviations

FdR: ferredoxin reductase
Fdx: ferredoxin
FjTAL: *F. johnsoniae* tyrosine ammonia lyase
PdR: *P. putida* putidaredoxin reductase
Pdx: *P. putida* putidaredoxin
PCNA: Proliferating Cell Nuclear Antigen = heterotrimeric DNA sliding clamp; used as tether
p. i.: post induction
PuR: *R. palustris* palustrisredoxin reductase
Pux: *R. palustris* palustrisredoxin
RgTAL: *R. glutinis* tyrosine ammonia lyase
TAL: tyrosine ammonia lyase

## Declarations

### Ethics approval and consent to participate

Not applicable.

### Consent for publication

Not applicable.

### Availability of data and materials

The datasets used and/or analyzed during the current study are available from the corresponding author on reasonable request.

### Authors’ contributions

K.H. conceived the study and wrote the manuscript with support and guidance by K.L.J.P. K.H. performed and analyzed the experiments.

## Acknowledgements

K.H. and K.L.J.P. are grateful to Dr. Stephen G. Bell (University of Adelaide, Australia) for plasmid pACYCDuet::PuR_Pux, and David Poberejsky for assistance with cloning. K.H. is grateful for the support by the Human Frontier Science Program (Grant Number LT000969/2016-L). This work was supported by the MIT Portugal Program (Grant Number 6937822).

