## Supporting Information for "Heterologous caffeic acid biosynthesis in *Escherichia coli* is affected by choice of tyrosine ammonia lyase and redox partners for bacterial Cytochrome P450"

### Sequences of codon optimized genes

FjTAL:

ATGAACACCATCAACGAATATCTGAGCCTGGAAGAATTTGAAGCCATTATCTTTGGCAATCAGAAAGTGACCATTAGTGATGTTGTTGTGAATCGCGTTAACGAGAGCTTTAACTTTCTGAAAGAATTTAGCGGCAACAAAGTGATCTATGGTGTGAATACCGGTTTTGGTCCGATGGCACAGTATCGTATTAAAGAAAGCGATCAGATTCAGCTGCAGTATAATCTGATTCGTAGCCATAGCAGCGGCACCGGTAAACCGCTGAGTCCGGTTTGTGCAAAAGCAGCAATTCTGGCACGTCTGAATACCCTGAGTCTGGGTAATAGCGGTGTTCATCCGAGCGTTATTAATCTGATGAGCGAACTGATCAACAAAGATATTACACCGCTGATTTTTGAACATGGTGGTGTTGGTGCAAGCGGTGATCTGGTTCAGCTGAGCCATCTGGCACTGGTTCTGATTGGTGAAGGTGAAGTTTTCTATAAAGGTGAACGTCGTCCGACACCGGAAGTTTTTGAAATTGAAGGTCTGAAACCGATCCAGGTGGAAATTCGCGAAGGTCTGGCCCTGATTAATGGCACCAGCGTTATGACCGGTATTGGTGTTGTTAATGTGTACCATGCAAAAAAACTGCTGGATTGGAGCCTGAAAAGCAGCTGTGCAATTAATGAACTGGTTCAGGCATATGATGATCACTTTAGCGCAGAACTGAATCAGACCAAACGTCATAAAGGTCAGCAAGAAATTGCACTGAAAATGCGTCAGAATCTGAGCGATAGCACCCTGATTCGCAAACGTGAAGATCATCTGTATAGCGGTGAAAACACCGAAGAAATCTTCAAAGAAAAAGTGCAAGAGTATTATAGCCTGCGTTGTGTTCCGCAGATTCTGGGTCCGGTTCTGGAAACCATTAACAATGTTGCAAGCATTCTGGAAGATGAATTTAACAGCGCAAACGATAACCCGATCATCGATGTTAAAAACCAGCATGTTTATCACGGTGGCAATTTTCATGGTGATTATATCAGCCTGGAAATGGATAAACTGAAAATCGTGATTACCAAACTGACCATGCTGGCAGAACGTCAGCTGAATTATCTGCTGAATAGCAAAATTAACGAACTGCTGCCTCCGTTTGTTAATCTGGGCACCCTGGGTTTTAACTTTGGTATGCAGGGTGTTCAGTTTACCGCAACCAGCACCACCGCAGAAAGCCAGATGCTGAGCAATCCGATGTATGTTCATAGCATTCCGAACAATAATGATAACCAGGATATTGTTAGCATGGGCACCAATAGCGCAGTTATTACCAGCAAAGTTATCGAAAATGCCTTTGAAGTTCTGGCCATTGAAATGATTACCATTGTTCAGGCGATTGATTATCTGGGCCAGAAAGATAAAATCAGCAGCGTTAGCAAAAAATGGTATGATGAAATCCGCAACATCATCCCGACCTTTAAAGAAGATCAGGTGATGTATCCGTTCGTGCAGAAAGTAAAAGACCACCTGATTAACAATtga

SeSam8:

ATGACCCAGGTTGTTGAACGTCAGGCAGATCGTCTGAGCAGCCGTGAATATCTGGCACGTGTTGTTCGTAGCGCAGGTTGGGATGCAGGTCTGACCAGCTGTACCGATGAAGAAATTGTTCGTATGGGTGCAAGCGCACGTACCATTGAAGAATATCTGAAAAGCGATAAACCGATCTATGGTCTGACCCAGGGTTTTGGTCCGCTGGTTCTGTTTGATGCAGATAGCGAACTGGAACAGGGTGGTAGCCTGATTAGCCATCTGGGCACCGGTCAGGGTGCACCGCTGGCACCGGAAGTTAGCCGTCTGATTCTGTGGCTGCGTATTCAGAATATGCGTAAAGGTTATAGCGCAGTTAGTCCGGTTTTTTGGCAGAAACTGGCAGATCTGTGGAATAAAGGTTTTACACCGGCAATTCCGCGTCATGGCACCGTTAGCGCAAGCGGTGATCTGCAGCCGCTGGCCCATGCAGCACTGGCATTTACCGGTGTTGGTGAAGCATGGACCCGTGATGCCGATGGTCGTTGGAGCACCGTTCCGGCAGTTGATGCACTGGCAGCCCTGGGTGCAGAACCGTTTGATTGGCCTGTTCGTGAAGCACTGGCCTTTGTTAATGGTACAGGTGCAAGCCTGGCAGTTGCAGTTCTGAATCATCGTAGTGCACTGCGTCTGGTTCGTGCATGTGCCGTTCTGAGCGCACGTCTGGCAACCCTGCTGGGTGCAAATCCGGAACATTATGATGTTGGTCATGGTGTTGCACGTGGTCAGGTTGGCCAGCTGACCGCAGCAGAATGGATTCGTCAGGGTCTGCCTCGTGGTATGGTTCGTGATGGTAGCCGTCCGCTGCAAGAACCGTATAGCCTGCGTTGTGCACCGCAGGTTCTGGGTGCGGTTCTGGATCAGCTGGATGGTGCCGGTGATGTTCTGGCACGCGAAGTTGATGGTTGTCAGGATAATCCGATTACCTATGAAGGTGAACTGCTGCACGGTGGTAATTTTCATGCAATGCCGGTTGGTTTTGCAAGCGATCAGATTGGTCTGGCAATGCATATGGCAGCATATCTGGCCGAACGTCAGCTGGGTCTGCTGGTTTCACCGGTTACCAATGGCGATCTGCCACCGATGCTGACACCGCGTGCAGGTCGTGGTGCAGGACTGGCAGGCGTTCAGATTAGCGCAACCAGCTTTGTTAGCCGTATTCGCCAGCTGGTTTTTCCGGCAAGCCTGACCACCCTGCCGACCAATGGTTGGAATCAGGATCATGTTCCGATGGCACTGAATGGTGCAAATAGCGTTTTTGAAGCCCTGGAACTGGGTTGGCTGACCGTGGGTAGCCTGGCCGTTGGTGTTGCCCAGCTGGCAGCAATGACAGGTCATGCAGCAGAAGGTGTTTGGGCTGAACTGGCAGGTATTTGTCCGCCTCTGGATGCCGATCGTCCACTGGGTGCCGAAGTTCGTGCAGCACGTGATCTGCTGAGTGCACATGCAGATCAGCTGCTGGTTGATGAAGCAGATGGTAAAGATTTTGGCtga

PCNA1-PdR:

ATGTTCAAAATCGTGTACCCGAACGCCAAAGATTTTTTCAGCTTTATTAACAGCATCACCAACGTGACCGATAGCATTATTCTGAACTTTACCGAAGATGGCATCTTTAGCCGTCATCTGACCGAAGATAAAGTTCTGATGGCAATTATGCGCATTCCGAAAGATGTTCTGAGCGAATATTCAATTGATAGCCCGACCAGCGTTAAACTGGATGTTAGCAGCGTGAAAAAAATCCTGAGCAAAGCAAGCAGCAAAAAAGCAACCATTGAACTGACCGAAACCGATAGCGGTCTGAAAATTATCATCCGTGATGAAAAAAGCGGTGCCAAAAGCACCATTTATATCAAAGCAGAAAAAGGCCAGGTTGAACAGCTGACAGAACCGAAAGTTAATCTGGCAGTGAATTTTACCACCGATGAAAGCGTTCTGAATGTTATTGCAGCAGATGTTACCCTGGTTGGTGAAGAAATGCGTATTAGCACCGAAGAGGACAAAATCAAAATTGAAGCCGGTGAAGAGGGTAAACGTTATGTTGCATTTCTGATGAAAGACAAGCCGCTGAAAGAACTGAGCATTGATACCAGCGCCAGCAGCAGCTATAGCGCAGAAATGTTTAAAGATGCAGTTAAAGGTCTGCGTGGTTTTAGCGCACCGACAATGGTGAGCTTTGGTGAAAATCTGCCGATGAAAATTGATGTTGAAGCAGTTAGCGGTGGCCACATGATTTTTTGGATTGCACCGCGTTTAGGTGGTGGTGGTAGCGGTGGTGGCGGTTCAATGAATGCAAATGATAATGTTGTTATCGTTGGCACCGGTCTGGCAGGCGTTGAAGTTGCATTTGGCCTGCGTGCAAGCGGTTGGGAAGGTAATATTCGTCTGGTGGGTGATGCAACCGTTATTCCGCATCATCTGCCTCCGCTGAGTAAAGCATATCTGGCAGGTAAAGCAACCGCAGAAAGCCTGTATCTGCGTACACCGGATGCCTATGCAGCACAGAATATTCAGCTGTTAGGTGGCACCCAGGTTACCGCAATTAATCGTGATCGTCAGCAGGTTATTCTGAGTGATGGTCGTGCACTGGATTATGATCGTCTGGTTCTGGCAACCGGTGGTCGTCCGCGTCCGCTGCCGGTTGCAAGTGGTGCAGTTGGTAAAGCCAATAACTTTCGTTATCTGCGCACCCTGGAAGATGCAGAATGTATTCGTCGTCAGCTGATTGCAGATAATCGCCTGGTTGTTATTGGTGGTGGCTATATTGGTCTGGAAGTTGCAGCAACCGCCATTAAAGCAAATATGCATGTGACCCTGCTGGATACCGCAGCACGTGTTCTGGAACGTGTTACCGCACCGCCTGTTAGCGCCTTTTATGAACATCTGCATCGTGAAGCGGGTGTTGATATTCGCACCGGTACACAGGTTTGTGGTTTTGAAATGAGCACCGATCAGCAGAAAGTTACCGCAGTTCTGTGTGAAGATGGTACACGTCTGCCTGCAGATCTGGTTATTGCCGGTATTGGCCTGATTCCGAATTGTGAACTGGCAAGCGCAGCAGGTCTGCAGGTTGATAATGGTATTGTTATTAACGAACACATGCAGACCAGCGATCCGCTGATTATGGCAGTTGGTGATTGTGCACGTTTTCATAGCCAGCTGTATGATCGTTGGGTTCGTATTGAAAGCGTGCCGAATGCACTGGAACAGGCACGTAAAATTGCAGCAATTCTGTGTGGCAAAGTTCCGCGTGATGAAGCAGCACCGTGGTTTTGGAGCGATCAGTATGAAATCGGCCTGAAAATGGTTGGTCTGAGTGAAGGTTATGATCGCATTATTGTTCGTGGTAGCCTGGCACAGCCGGATTTTTCAGTTTTTTATCTGCAGGGTGATCGTGTGCTGGCAGTTGATACCGTTAATCGTCCGGTTGAATTTAATCAGAGCAAGCAGATTATTACCGATCGTCTGCCGGTGGAACCGAACCTGCTGGGTGATGAAAGTGTTCCTCTGAAAGAAATTATTGCCGCAGCAAAAGCAGAACTGAGTAGCGCATAA

### Supplementary Tables

Table S1: Comparison of select tyrosine ammonia lyase enzymes characterized in other studies.

| Enzyme | K_m_ [mM] | k_cat_ [s^-1^] | k_cat_/K_m_ [mM^-1^ s^-1^] | Ratio TAL:PAL activity | Reference |
| --- | --- | --- | --- | --- | --- |
| RgTAL (enzyme purified by anion exchange chromatography, ammonium sulfate precipitation and hydrophobic interaction chromatography; assay performed at pH 8.5; T=25°C) | 615 | 0.53 | 8.6×10^-4^ | 10 | [1] |
| RgTAL (enzyme purified by anion exchange chromatography, ammonium sulfate precipitation and hydrophobic interaction chromatography; assay performed at pH 9.5; T=25°C) | 67.7 | 0.93 | 1.37x10^-2^ | 10 | [1] |
| FjTAL (enzyme purified by metal affinity chromatography; assay performed at pH 9.5; T=30°C) | 6.7x10^-3^ | 0.023 | 2.99 | 2400 | [2] |
| SeSam8 (enzyme purified by metal affinity chromatography; assay performed at pH 9.5; T=30°C) | 4.7x10^-3^ | 0.015 | 3.05 | 1200 | [2] |
| RgTAL (enzyme purified by metal affinity chromatography; assay performed at pH 8.5; T=40°C) | 0.38 | 114 | 298 | n.d. | [3] |

Supplementary Table 2: Plasmids generated for this study including cloning strategies used.

| **plasmid name [backbone::MCSI_MCSII]** | **cloning method** | **Primers (binding sequence in lowercase, overhang in uppercase letters)** | **source** |
| --- | --- | --- | --- |
| IR54 pKVS45::PdR-Pux operon | / | / | [4] |
| IR64 pCDFduet::_6His-CYP199A2 F185L NΔ7 | / | / | [4] |
| c22 pRSFduet::6His-RgTAL | Golden Gate Assembly enzyme: SapI template: IR66 | insert MCSI fwd CGTCAACGCTCTTCCtccgcgtccgacctcgca insert MCSI rev CGTCAACGCTCTTCCcttatgccagcattttcagcagc backbone fwd CGTCAACGCTCTTCCaagcggccgcataatgctta backbone rev CGTCAACGCTCTTCCggagccatttggcgcgccgagctcga | This study |
| c25 pCDFduet::_PCNA3-CYP199A2 F185L NΔ7 | Golden Gate Assembly enzyme: BsaI templates: pHSG-PCNA3 and IR66 | PCNA3 fwd CGTCAACGGTCTCGACATatgatatatcttaaatcttttgaaaggaatataag PCNA3 rev CGTCAACGGTCTCGGCATagatccaccaacttttggagc CYP199A2 F185L NΔ7 fwd CGTCAACGGTCTCGatgccggttacgacgccg CYP199A2 F185L NΔ7 rev CGTCAACGGTCTCGGACAtcaggccggggtcagttg backbone fwd CGTCAACGGTCTCGtgtcttcggtaccctcgag backbone rev CGTCAACGGTCTCGatgtatatctccttcttatacttaactaa | This study |
| c28 pETduet::6His-PCNA2-Pux_PCNA1-PdR (opt) | classic cloning  MCSII: NdeI/KpnI (insert: gblocks), backbone: c58 | insert MCS II fwd GATCTACATatgttcaaaatcgtgtacccgaa insert MCSII rev GATCTAggtaccactagtatttatgcgctac | This study |
| c50 pETduet::6His-Pux_PdR (opt) | classic cloning  MCSI: SacI/NotI (template: IR54),  MCSII: NdeI/EcoRV (template: gblocks) | insert MCSI fwd GATCGAGCTCAatgcccagtatcacgttcatt insert MCSI rev GATCgcggccgcaagcttgtcg insert MCSII fwd GATCATCATatgaatgcaaatgataatgttgttatcgttg insert MCSII rev GATCTAGGTACCactagtatttatgcgctac | This study |
| c62 pETduet::6His-Pux_PuR | classic cloning  MCSI: SacI/NotI (template: IR54),  MCSII: NdeI/KpnI (template: pACYCduet::PuR_Pux) | insert MCSI fwd GATCGAGCTCAatgcccagtatcacgttcatt insert MCSI rev GATCgcggccgcaagcttgtcg insert MCSII fwd GATCTACATatggacgacacggtcttgattg insert MCSII rev GATCGGTACCGAAGACAttacgccgccgccttcttc | This study |
| c63 pETduet::6His-PCNA2-Pux_PCNA1-PuR | restriction digest/ligation, enzymes: MCS2, NdeI/KpnI (insert: c64), backbone: c28 | / | This study |
| c71 pRSFduet::6His-FjTAL | classic cloning  enzymes: BamHI/ NotI (template: gblocks) | insert MCSI fwd GATCAggatccgagcagcggc insert MCSI rev GATCaagcggccgcaagctttca | This study |
| c72 pRSFduet::6His-SeSam8 | classic cloning  enzymes: BamHI/ NotI (template: gblocks) | insert MCSI fwd GATCAggatccgagcagcggc insert MCSI rev GATCaagcggccgcaagctttca | This study |
| c84 pCDFduet::6His-Pux_6His-CYP199A2F185L NΔ7 | MCSI: classic cloning, enzymes: SacI/NotI (template: c50); MCSII: restriction digest/ligation, NdeI/AvrII (insert: IR64) | insert MCSI fwd GATCGAGCTCAatgcccagtatcacgttcatt insert MCSI rev GATCgcggccgcaagcttgtcg | This study |
| c86 pETduet::6His-Pdx_PdR (opt) | classic cloning  enzymes: MCSI, SacI/HindIII, template: *Pseudomonas putida* gDNA; backbone:c50 | insert MCSI fwd GATCATGAGCTCAatgtctaaagtagtgtatgtgtcacatg insert MCSI rev GATCATAAGCTTGTCGACttaccattgcctatcgggaacatc | This study |
| c88 pETduet::6His-PCNA2-Pdx_PCNA1-PdR (opt) | classic cloning  enzymes: MCSI, BseRI/SalI, template: *Pseudomonas putida* gDNA; backbone: c28 | insert MCSI fwd GATCATCCTCCACCGCCTCCTCCACCGCCACCACCGCCGCCTCCAC CTCCACCGCCGCCCGGTatgtctaaagtagtgtatgtgtcacatg insert MCSI rev GATCATAAGCTTGTCGACttaccattgcctatcgggaacatc | This study |
| c96 pCDFduet::_PCNA1-GGS-CYP199A2 F185L NΔ7 | Round-the-Horn PCR to shorten peptide linker (template:c77) | fwd cggttcaatgccggttacg rev cctaaacgcggtgcaatccaaa | This study |
| c97 pETduet::6His-PCNA2-Pux_PCNA3-(GGGS)2-PdR (opt) | Round-the-Horn PCR to expand peptide linker (template: c80) | fwd AGCGGTGGTggtggatctatgaatgcaaatgataatgttg rev ACCACCACCACCaacttttggagctaataaataagtaactttccc | This study |
| c98 pETduet::6His-PCNA2-Pux_PCNA3-(GGGS)2-PuR | Round-the-Horn PCR to expand peptide linker (template: c81) | fwd AGCGGTGGTggtggatctatggacgacacg rev ACCACCACCACCaacttttggagctaataaataagtaactttccc | This study |
| c106 pETduet::6His-PCNA2-Pdx_PCNA3-GGS-PdR (opt) | restriction digest/ligation, enzymes: MCS2, NdeI/KpnI (insert: c97), backbone: c88 | / | This study |
| **PCR templates and additional plasmids** |  |  |  |
| IR66 pCDFduet::RgTAL_6His-CYP199A2 F185L NΔ7 | / | / | [4] |
| c55 pCDFDuet::_CYP199A2F185L NΔ7 (without His-Tag) | Golden Gate Assembly  enzyme: BsaI  template: IR66 | insert fwd CGTCAACGGTCTCGacatatgccggttacgacgccg  insert rev CGTCAACGGTCTCGgacatcaggccggggtcagttg  backbone fwd CGTCAACGGTCTCGacatatgccggttacgacgccg  backbone rev CGTCAACGGTCTCGacatatgccggttacgacgccg | This study |
| c58 pETduet::6His-PCNA2-Pux_ | MCSI: Golden Gate Assembly  enzyme: BsaI  templates: pHSG-PCNA2 (two-step PCR) and IR54 | PCNA2 fwd CGTCAACGGTCTCCCTCAatgaaagctaaggtaattgacgctg PCNA2 rev 1 CGTCAACGGTCTCCTGGCGGTGGAGGAGGCGGTGGAGGTGGAGGG CTGCCGCCACCgtctgcccttggtgcaatg PCNA2 rev 2 CGTCAACGGTCTCCTGGCGGTGGAGGAGGCGGTG Pux fwd CGTCAACGGTCTCCGCCACCACCGCCGCCTCCACCTCCACCGCCGCCCGGT atgcccagtatcacgttcattc Pux rev CGTCAACGGTCTCGCGACtcagacctgacgatccggaatc backbone fwd CGTCAACGGTCTCGgtcgacaagcttgcggccgc backbone rev CGTCAACGGTCTCGtgagctcgaattcggatcctggctgtggtga | This study |
| c64 pETDuet::_PCNA1-PuR | Golden Gate Assembly  enzyme: BsaI  templates: c28 and pACYC::PuR_Pux | PCNA1 fwd CGTCAACGGTCTCGACATtttaagattgtttaccctaatgcaaaagac PCNA1 rev CGTCAACGGTCTCGactcccgccgccaccagaac PuR fwd CGTCAACGGTCTCGGAGTatggacgacacggtcttgattg PuR rev CGTCAACGGTCTCGGACAttacgccgccgccttcttc backbone fwd CGTCAACGGTCTCGtgtcttcggtaccctcgag backbone rev GGTAAACAATCTTAAAtgagctcgaattcggatcctgg | This study |
| c77 pCDFduet::_PCNA1-(GGGS)2-CYP199A2 F185L NΔ7 | CPEC  template: c28, backbone: c55 | insert fwd tataagaaggagatatacatATGTTCAAAATCGTGTACCCG insert and CPEC rev GACGGCGTCGTAACCGGCATtgaaccgccaccaccgctac CPEC fwd GTAGCGGTGGTGGCGGTTCAatgccggttacgacgccgtc | This study |
| c78 pETduet::6His-PCNA2-Pux_PdR | classic cloning, enzymes: MCSII NdeI/KpnI (template: c28, backbone: c28) | insert MCSII fwd GATCATCATatgaatgcaaatgataatgttgttatcgttg insert MCSII rev GATCTAGGTACcactagtatttatgcgctac | This study |
| c79 pETduet::6His-PCNA2-Pux_PuR | classic cloning, enzymes: MCSII NdeI/KpnI (template: c62, backbone: c28) | insert MCSII fwd GATCTACATatggacgacacggtcttgattg insert MCSII rev GATCGGTACCGAAGACattacgccgccgccttcttc | This study |
| c80 pETduet::6His-PCNA2-Pux_PCNA3-PdR (GGS) | CPEC  template: c25 , backbone: c78 | insert fwd AGTATAAGAAGGAGATATACATatgatatatcttaaatcttttgaaaggaatataag attga insert and CPEC rev ACATTATCATTTGCATTCATagatccaccaacttttggag CPEC fwd CTCCAAAAGTTGGTGGATCTatgaatgcaaatgataatgttgtt | This study |
| c81 pETduet::6His-PCNA2-Pux_PCNA3-PuR (GGS) | CPEC  template: c25 , backbone: c79 | insert fwd AGTATAAGAAGGAGATATACATatgatatatcttaaatcttttgaaaggaatataag attga insert and CPEC rev ATCAAGACCGTGTCGTCCATagatccaccaacttttggag CPEC fwd CTCCAAAAGTTGGTGGATCTatggacgacacggtcttgat | This study |
| pACYCduet::PuR_Pux | / | / | Dr. Stephen G. Bell |
| pHSG-PCNA2 | / | / | [5] |
| pHSG-PCNA3 | / | / | [5] |

### Supplementary Figures

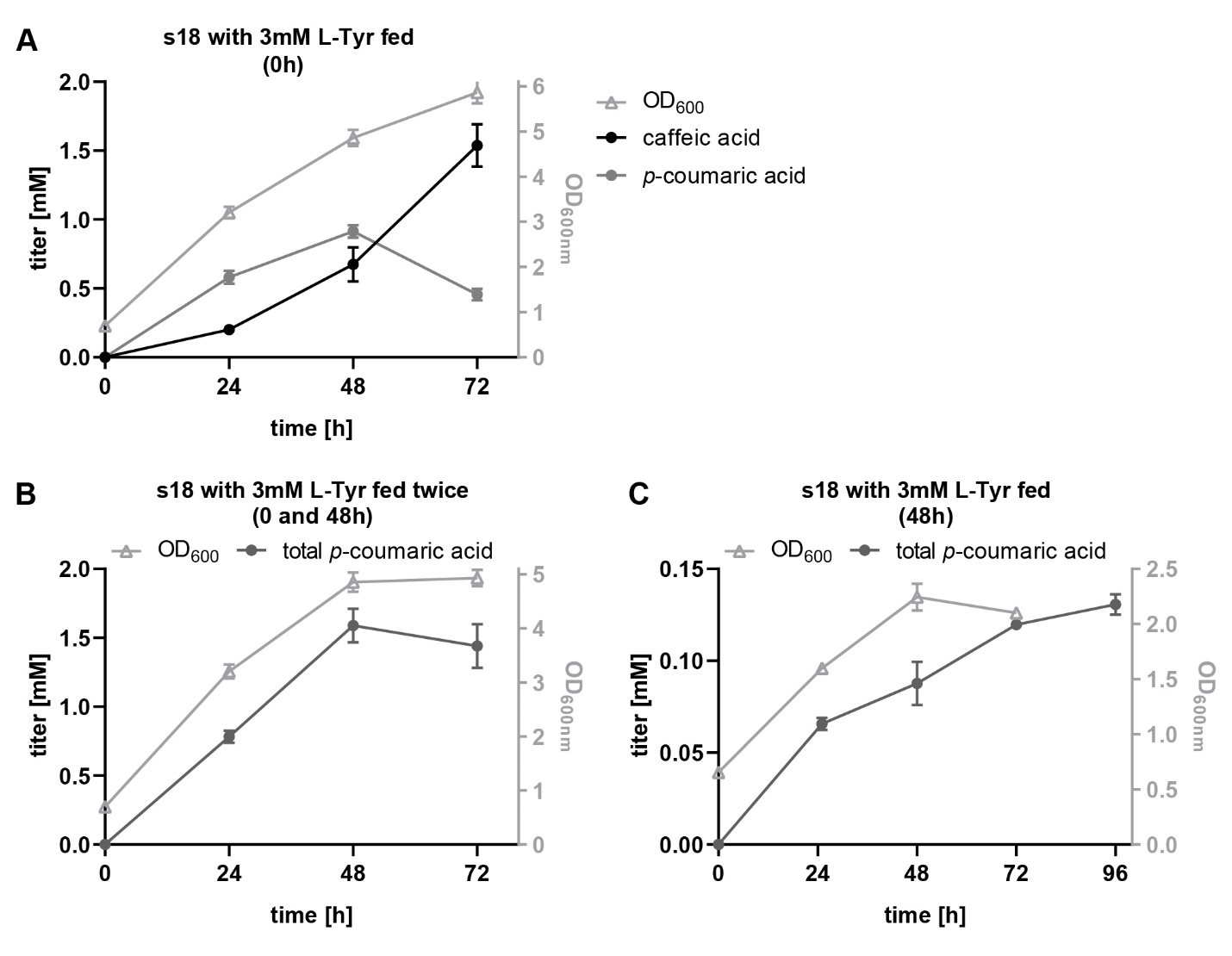

Figure S1: Time course experiments with strain s18 expressing the Pux/PuR redox system with a doubled gene dose of pux under different L-tyrosine feeding strategies and 4% (w/v) of glucose. Feeding of 3 mM L-tyrosine at t=0h p.i. (A), at t=0h p.i. and t=48h p.i. (B) and at t=48h p.i.(C). Total p-coumaric acid titers plotted in panels B and C were calculated by summing up the measured titers of p-coumaric and caffeic acid. In these experiments, no distinct increase in titers was observed after the addition of L-tyrosine at 48h p.i.. This indicates that the additional tyrosine could not be converted to coumaric or caffeic acid, most likely because the tyrosine ammonia lyase enzyme was inactive or denatured. (error bars=standard deviation of biological replicates, n=3).
